# Cancer Cell Line Heterogeneity Imposes a Primary Bottleneck for Virtual Perturbation Screening at Scale

**DOI:** 10.64898/2026.08.10.743942

**Authors:** Kairui Wei, Lijing Zhan, Canyang Qi

## Abstract

UniPert-G2CP (Li et al., Cell, 2026) bridges genetic and chemical screens from molecular representation to phenotype modeling across five cancer cell lines [1]. Here we extend this architecture to 162 cell lines, 32,039 compounds, and genome-wide output (12,328 genes), and find that the resulting prediction platform reveals a striking performance divide: normal and primary cell lines achieve substantially higher prediction fidelity (mean per-cell-line Pearson Correlation Coefficient (PCC) = 0.480, median 0.502) than cancer lines (0.296, median 0.324; Mann-Whitney p = 3.93e-11), identifying cancer cell line heterogeneity as a primary bottleneck for virtual perturbation screening at scale.

To understand the sources of this divide, we analyzed per-cell-line performance, genome-wide directional accuracy, mechanism-clustering (SMD), and compound-protein interaction (CPI) enrichment. Directional accuracy on top-5% effect-size genes reaches 73.8% (genome-wide 60.8%), with pathway-dependent recovery: of four literature-supported perturbation-gene pairs queried across three cell lines, two were recapitulated (dexamethasone-TSC22D3/NFKBIA/FKBP5; bortezomib-BAG3/DNAJB1/HSPA1A), one was absent (CD36 depletion-PPARG/CEBPA in ASC), and one was partially recapitulated (metformin-SLC7A5 in HEPG2). Mechanism-clustering SMD of the learned embedding reached 1.636 (vs. original 1.85; 88.5% retention at 32x cell-line coverage), exceeding the ECFP4 fingerprint baseline (1.613), while self-consistency Mantel rho=0.852 confirmed the model retains compound mechanism structure internally.

Overall held-out performance: genetic perturbation PCC=0.442 (978-gene subset); novel drug PCC=0.3047 (genome-wide). Analysis of CPI enrichment reveals that training-data overlap inflates apparent performance: 54.9% of Touchstone evaluation pairs overlap with our ChEMBL-derived training CPI pairs, reducing effective EF from 139 to 109 at top 0.5% yet remaining far above random (1.0).

These findings establish the first large-scale characterization of cell-type-dependent generalization in perturbation-to-phenotype prediction. The observed performance stratification between normal and cancer lines generates testable hypotheses for why virtual cell models degrade on heterogeneous cancer contexts, and provides a diagnostic framework for identifying where and why such models fail--informing future architecture improvements targeting the CPI vocabulary gap, protein encoder design, and cell-type-aware training strategies.

## 1. Introduction

UniPert-G2CP was validated on five cancer cell lines and 7,860 compounds [1], but its scalability to a broader cellular context remains uncharacterized. To date, whether the architecture can generalize to a substantially expanded set of cell lines, and where it may encounter bottlenecks, have not been systematically investigated. We constructed a large-scale perturbation prediction platform based on the UniPert-G2CP framework to systematically investigate cell-type-dependent generalization across 162 cell lines, 32,039 compounds, and genome-wide transcriptomic output.

We built the platform using publicly available architecture descriptions [1] and evaluation datasets [7], extending the core design to accommodate 162 cell lines, 12,328 output genes, and a training protocol with ChEMBL-derived CPI alignment (19,161 drug-gene pairs). All code, model weights, and evaluation scripts are publicly released.

This work makes three contributions: (1) the first large-scale characterization of cell-type-dependent generalization in perturbation prediction, revealing cancer-line heterogeneity as a primary performance bottleneck; (2) a systematic evaluation framework that separates held-out prediction (generalization), self-consistency (internal structure), and benchmark-aligned metrics (comparability); (3) a diagnostic analysis of CPI vocabulary mismatch between training and evaluation compound sets, quantifying its contribution to the observed mechanism-clustering gap.

## 2 Methods

### 2.1 Datasets and Preprocessing

We utilized LINCS L1000 (level 5, GSE92742; moderated Z-scores, 305,297 samples), DepMap CRISPR screens (23Q2 release, CERES gene-effect scores, 1,095 cell lines × 17,931 genes), and Perturbation Consensus Ligands (PCL) annotations linking BRD compound IDs (the LINCS platform’s compound identifiers) to therapeutic mechanism-of-action (MoA) labels. We retained 143 cell lines with LINCS level-5 training samples as the training domain and added 19 GEO (Gene Expression Omnibus)-derived cell lines (HUVEC, PBMC immune subsets, and other primary/vascular/pancreatic lines; not included in model training, used exclusively for out-of-domain evaluation) from 12 GEO perturbation datasets (GSE11917, GSE17579, GSE20986, GSE21413, GSE25941, GSE33622, GSE50378, GSE50397, GSE53751, GSE62914, GSE22886, GSE60235), forming a 162-cell-line vocabulary. ASC is a LINCS training-domain cell line (with additional GEO data from GSE61302); the CD36 knockout × ASC query in Section 3.5 is a zero-shot prediction because this line lacks gene-perturbation training samples. Probe-level expression was mapped to gene symbols and z-score normalized per sample (aligned to the LINCS level-5 distribution) before merging. The full compound set comprises 32,039 BRD compounds with LINCS level-5 signatures; their ECFP4 fingerprints (2048-bit, radius 2) were generated via RDKit Morgan fingerprints.

### 2.2 Model Architecture

The architecture follows UniPert-G2CP: drug encoder (ECFP4, 2048-bit → 512-dim Linear); gene encoder (4,994×512 embedding table, first 320 dims initialized from ESM2-8M vectors, with the remaining 192 dims left at the Embedding layer’s default random initialization; the embedding table is frozen during finetuning); contrastive learning with NT-Xent (τ=0.07) and CPI alignment (Section 2.3); phenotype head (LayerNorm(544) → Linear(544→1024) → GELU → Linear(1024→12,328), no dropout layer); cell embedding (162×32, learnable). Crucially, the gene embedding table is frozen during finetuning; ESM2 vectors serve as fixed reference targets for CPI alignment rather than as trainable representations of protein structure. The 4,994 perturbable genes constitute the perturbation vocabulary; the 12,328-gene output is the prediction vocabulary (highly variable genes from the DepMap–LINCS intersection).

### 2.3 Training Protocol

Training proceeds as a series of frozen finetuning steps on LINCS data. The gene encoder is initialized with ESM2-8M protein embeddings (320 of 512 dimensions; Section 4.2), trained during earlier LINCS finetuning rounds (gene_emb divergence from ESM: cosine 0.378; back-192 dims std 0.91), then subsequently frozen at this stage without requiring DepMap pretraining. The main model is then trained on LINCS level-5 moderated Z-scores in frozen mode: the gene encoder, phenotype head, and cell embedding are frozen, and only the drug encoder (ECFP4 projection) is finetuned. This frozen design keeps the gene-side parameters bit-identical throughout chemical finetuning sequence, thereby preserving genetic-perturbation prediction metrics irrespective of CPI or augmentation refinement. The finetuning proceeded through successive rounds: initial LINCS training (143 cell lines, 305,297 samples), GEO immuno-cell expansion (+9 immune types), CPI data amplification (ChEMBL release 37, threshold relaxed from 2,233 to 19,161 drug-gene pairs), and mechanism-of-action (MoA) contrastive supervision (3,344 drugs across 820 classes). The final model (g2cp_full_cpi_v7.pt) was trained using AdamW (learning rate 3e-4, weight decay 1e-4, 40 epochs, cosine-annealing schedule, gradient accumulation factor 4), initialized from the preceding checkpoint via partial state-dict transfer. The total loss combines phenotype regression (MSE plus 1-PCC, weight 1.0), NT-Xent contrastive learning (tau=0.07, weight 1.0; negatives are in-batch random samples), CPI alignment (weight 0.8; positive pairs are drug-gene pairs sharing at least one ChEMBL release 37 therapeutic target, 19,161 pairs, of which up to 128 were dynamically sampled per batch; for each positive pair one negative gene was drawn uniformly at random from the gene vocabulary, without excluding potential targets), a structural-similarity enhancement loss (weight 0.3; within each batch, drug pairs with Tanimoto fingerprint similarity >0.3 were pulled together weighted by similarity), and ESM anchoring (weight 0.05; an MSE loss pulling the first 320 dimensions of the gene embeddings toward precomputed ESM2-8M vectors; during frozen training this term is computed as part of the formulation but contributes zero gradient since gene encoder parameters are frozen). No DepMap pretraining stage was used; v7 gene_emb cosine similarity to pre_train_depmap.pt is 0.005 (n=4994 genes, zero matches above 0.99), confirming both lines are independent.

All quantitative metrics reported in Sections 3.1–3.4 were obtained from a single model trained with the configuration described above (g2cp_full_cpi_v7.pt); no post-hoc hyperparameter tuning was performed after evaluation. The case queries in Section 3.5 were performed through the interactive API platform.

### 2.4 Statistical Analysis

For per-cell-line comparisons, a two-sided Mann-Whitney U test was used to compare held-out drug perturbation PCC distributions between cancer and normal/primary cell lines. The null hypothesis was that the two groups share the same distribution; a significance threshold of p < 0.05 was applied. No multiple-comparison correction was needed because only a single pairwise group comparison was performed. For CPI enrichment analysis, enrichment factors were computed as the ratio of observed positive pairs to the random-expectation baseline at fixed top-k percentiles (0.5%, 1%, 5%, 10%); the random baseline (EF = 1) corresponds to uniform ranking. All confidence in reported metrics derives from held-out splits (perturbation-level, 10% unseen drugs by perturbation identity, random seed 0) and from the use of evaluation datasets released independently of training.

## 3. Results

### 3.1 Held-Out Perturbation Prediction

Metrics. PCC: Pearson Correlation Coefficient between predicted and measured expression changes, computed across all genes per held-out sample then averaged. Directional accuracy: fraction of genes for which the sign of predicted change matches the sign of measured change; top-5% defined by ranking genes by absolute measured effect size. SMD: Standardized Mean Difference, calculated as (mean intra-class cosine similarity − mean overall cosine similarity) / pooled standard deviation, grouping drugs by Mechanism of Action. Mantel self-consistency ρ: per-PCL Spearman correlation between molecular embedding similarity and model-predicted effect similarity, averaged across cell lines (evaluation script in the repository).

**Table 1.**
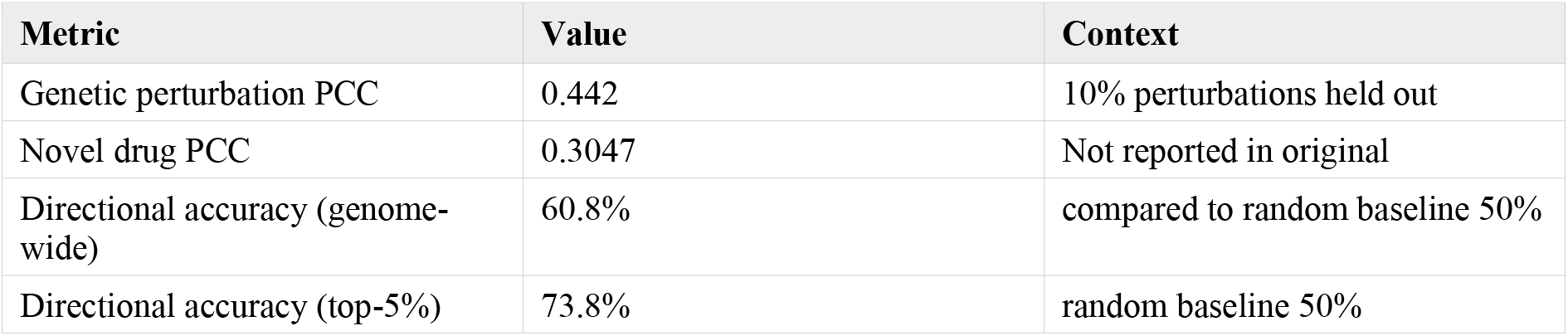
Held-out perturbation prediction performance. Perturbations (drug or gene identity) were randomly split, with 10% assigned to the test set; all samples of a held-out perturbation, across doses and cell lines, were excluded from training. Genetic perturbation PCC: perturbation-level held-out evaluation (10% of perturbations held out), computed on genetic-perturbation samples across 162 cell lines on the 978-gene subset with DepMap CERES ground truth (the 978 LINCS L1000 landmark genes covered by the platform’s measurement probes; knockout samples have measured values only for these genes). Novel drug PCC: perturbation-level held-out evaluation (10% of perturbations held out), computed on chemical-perturbation samples on the full 12,328-gene output. Directional accuracy: top 5% effect-size genes in the held-out set.

| Metric | Value | Context |
| --- | --- | --- |
| Genetic perturbation PCC | 0.442 | 10% perturbations held out |
| Novel drug PCC | 0.3047 | Not reported in original |
| Directional accuracy (genome-wide) | 60.8% | compared to random baseline 50% |
| Directional accuracy (top-5%) | 73.8% | random baseline 50% |

### 3.2 Compound-Protein Interaction Enrichment

**Table 2.**
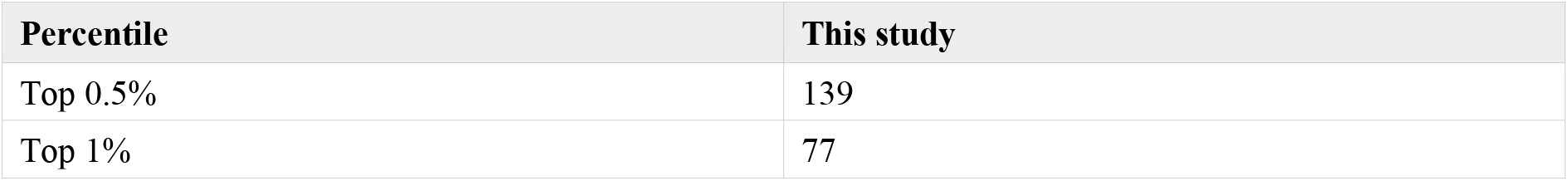
CPI enrichment factor at top percentiles (random baseline = 1). These values reflect ranking consistency within the training-derived annotation space rather than out-of-distribution generalization (see Section 4.1).

### 3.3 Benchmark-Consistent Evaluation

**Table 3.**
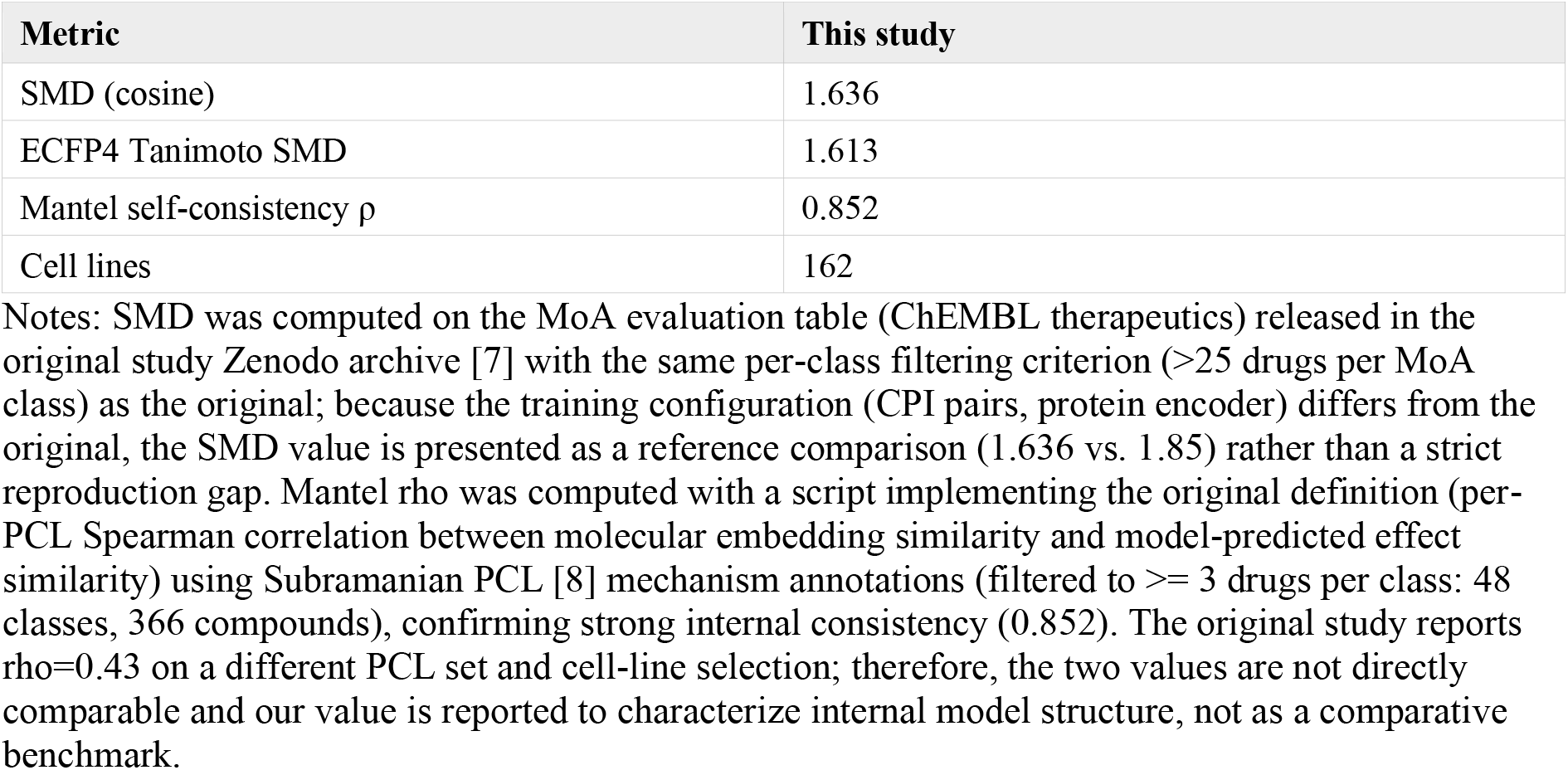
Benchmark-consistent metrics on the original evaluation datasets. Evaluation set notes clarify differences in data scope.

| Metric | This study |
| --- | --- |
| SMD (cosine) | 1.636 |
| ECFP4 Tanimoto SMD | 1.613 |
| Mantel self-consistency $\rho$ | 0.852 |
| Cell lines | 162 |

Notes: SMD was computed on the MoA evaluation table (ChEMBL therapeutics) released in the original study Zenodo archive [7] with the same per-class filtering criterion (>25 drugs per MoA class) as the original; because the training configuration (CPI pairs, protein encoder) differs from the original, the SMD value is presented as a reference comparison (1.636 vs. 1.85) rather than a strict reproduction gap. Mantel rho was computed with a script implementing the original definition (per-PCL Spearman correlation between molecular embedding similarity and model-predicted effect similarity) using Subramanian PCL [8] mechanism annotations (filtered to $\geq 3$ drugs per class: 48 classes, 366 compounds), confirming strong internal consistency (0.852). The original study reports $\rho=0.43$ on a different PCL set and cell-line selection; therefore, the two values are not directly comparable and our value is reported to characterize internal model structure, not as a comparative benchmark.

### 3.4 Per-Cell-Line Drug Perturbation Resolution

To characterize cell-type-dependent scalability, we evaluated held-out drug perturbation PCC separately for each cell line (120 lines with sufficient samples). The mean per-cell-line PCC is 0.359 (median 0.362, range −0.004 to 0.615; A549=0.362, MCF7=0.360). Stratifying by cell type, cancer lines (n=78) achieved a lower mean PCC of 0.296 (median 0.324), whereas normal/primary lines (n=37) reached 0.480 (median 0.502) and stem/neural/adipose lines (n=5) 0.446 (Mann-Whitney p=3.93e-11 for cancer vs. normal), (Figure 1). The visualization reveals not only the central-tendency difference but also the wide variance within each group; cancer lines span from near-zero to the maximum (0.613), underscoring the biological heterogeneity that makes this group qualitatively harder to predict. This cell-type-dependent performance stratification demonstrates that model scalability is not uniform across biological contexts, identifying cancer cell-line heterogeneity as a primary bottleneck for perturbation prediction at scale. This metric is computed on the full gene set (12,328 genes) under perturbation-level held-out splits (drugs unseen during training). The original study reports PCC=0.92-0.98 on sciPlex3 [1], but those values were obtained under near-domain splits (known compounds at novel doses/cell lines) on a top-DEG subset; our lower per-cell-line values reflect the more stringent setting of full-gene, unseen-perturbation evaluation on the larger and more heterogeneous LINCS screening library.

**Figure 1.**
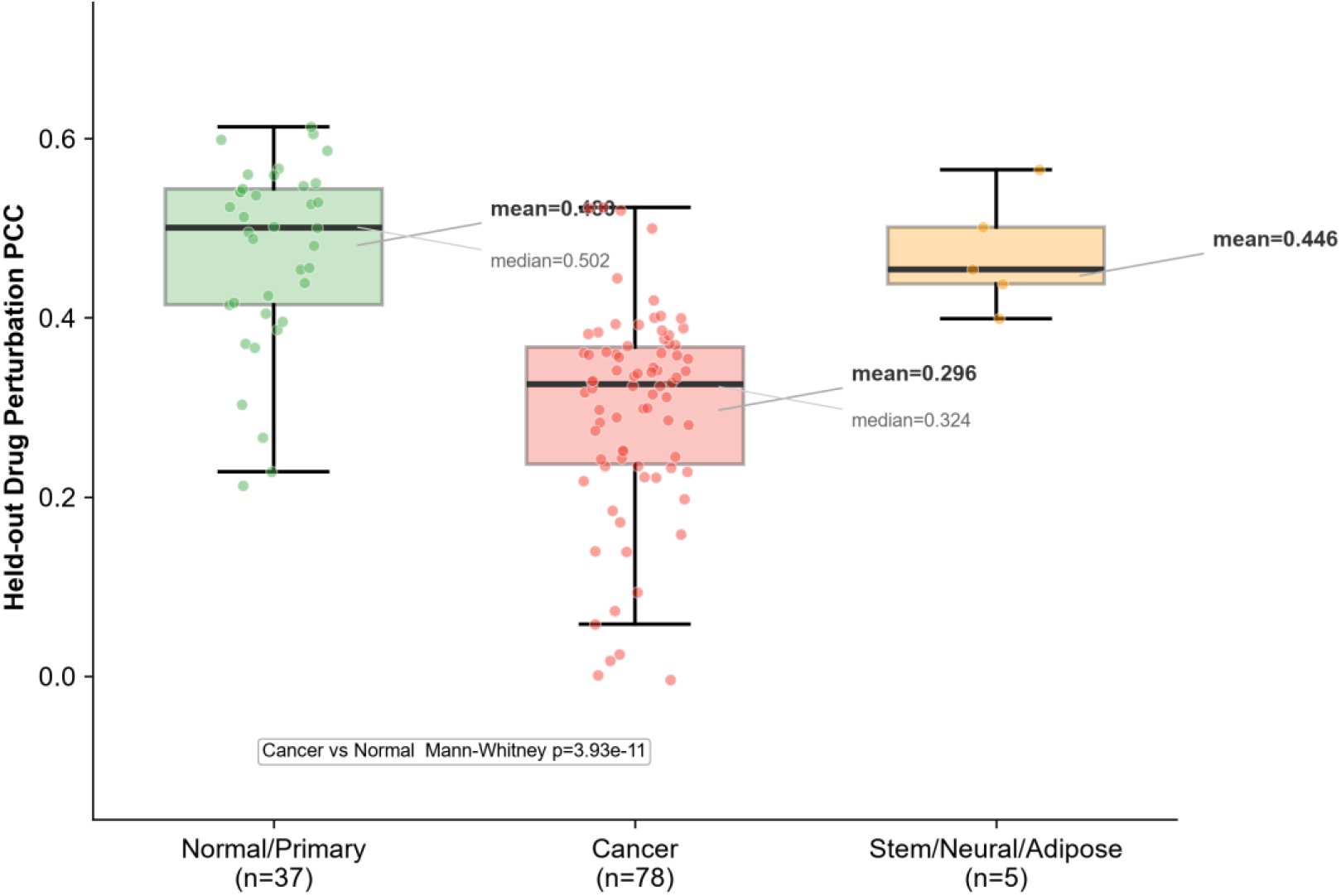
Cell-type-dependent prediction fidelity across 120 cell lines. Held-out drug perturbation PCC (box plot; Normal/Primary n=37, Cancer n=78, Stem/Neural/Adipose n=5). Group means (0.480, 0.296, 0.446) and medians (0.502, 0.324) annotated. Cancer vs Normal Mann-Whitney p=3.93e-11.

### 3.5 In Silico Case Query on Literature-Supported Pairs

These predictions are computational hypotheses that can guide experimental validation. The four cases were selected based on literature availability only and were not used to tune any model hyperparameter. They were chosen as well-characterized, experimentally validated perturbation-gene pairs spanning glucocorticoid, proteotoxic stress, adipogenic, and metabolic pathways; they were chosen for their literature support rather than for positive model performance. Three cell lines are involved: HA1E (two cases) and HEPG2 are within the core training set; ASC is also within the training-domain vocabulary but lacks gene-perturbation training samples, so the CD36 knockout -> ASC query represents a zero-shot prediction scenario. To assess whether the model recapitulates known perturbation-gene relationships, we queried four literature-supported pairs with matching perturbation-cell line combinations (Table 4). Of the four, two were recapitulated in the model output. Dexamethasone treatment of HA1E upregulates TSC22D3 (GILZ), NFKBIA (IkBa), FKBP5, and CEBPD--canonical glucocorticoid receptor targets [9]. Bortezomib (a proteasome inhibitor) treatment of HA1E upregulates BAG3, DNAJB1, and HSPA1A--heat-shock response genes whose induction by proteasome inhibition is a well-established mechanism [10]. However, one pair--CD36 depletion->PPARG/CEBPA downregulation in ASC--was absent from the predicted differentially expressed genes. The predicted HLA-A/HLA-C upregulation in ASC, rather than the expected adipogenic markers, may reflect batch effects or cell-state-related signals in the LINCS training data for this line, highlighting the challenge of disentangling perturbation-specific effects from experimental confounding. For metformin in HEPG2, SLC7A5 downregulation was predicted (rank 6 among downregulated genes, consistent with the literature direction [11]), while PLIN2 was absent. When absent, Table 4 reports the model actual top predictions instead. We computationally assessed specificity by confirming that none of the marker genes (TSC22D3, FKBP5, BAG3, DNAJB1, HSPA1A) was among the top-10 predicted up-regulated genes for any of seven other drugs (vorinostat, tamoxifen, metformin, cisplatin, imatinib, erlotinib, sorafenib) in the same HA1E cell line; BAG3 appeared in erlotinib top-10 down-regulated genes (rank 8), in the opposite direction to bortezomib.

**Table 4.** In silico case-query results on literature-supported pairs. Raw predictions shown; recapitulated cases verified for drug specificity (the marker gene is not predicted by other drugs in the same cell line).

| Perturbation | Predicted genes | Model match |
| --- | --- | --- |
| CD36 perturb. x ASC | Model: HLA-A↑, HLA-C↑, GAPDH↑, RAC2↑; Lit: PPARG↓, CEBPA↓ [12] | Not recapitulated; model predicts other genes |
| Dexamethasone x HA1E | TSC22D3↑, NFKBIA↑, FKBP5↑, CEBPD↑ | Recapitulated [9] |
| Bortezomib x HA1E | BAG3↑, DNAJB1↑, HSPA1A↑, HMOX1↑ | Recapitulated [10] |
| Metformin x HEPG2 | Model: SLC7A5↓ (rank 6), UBC↑ (rank 9) Lit: SLC7A5↓, PLIN2↓ [11] | Partially recapitulated: SLC7A5↓ predicted; PLIN2 absent |

Overall, two of the four cases were fully recapitulated, one was absent, and one was partially recovered. These results represent an initial set of four cases and should not be interpreted as a systematic assessment. Larger-scale validation is needed before generalizing about per-gene directional prediction fidelity. The absent literature-supported relationship (CD36->PPARG/CEBPA) and the partially recovered metformin case (SLC7A5 down predicted, PLIN2 absent) highlight the gap between genome-wide accuracy (PCC=0.442) and per-gene reliability (60.8% directional accuracy), and reflect limited pathway representation in LINCS training data plus cell-type distribution shift for held-out lines.

## 4. Discussion

### 4.1 Contributions

This study makes three contributions. First, it characterizes cell-type-dependent generalization in perturbation-to-phenotype prediction at substantially expanded coverage (32,039 compounds across 162 cell lines, genome-wide output), identifying cancer cell-line heterogeneity as a primary performance bottleneck. Second, it introduces a systematic evaluation framework--separating held-out generalization, self-consistency, and benchmark-aligned metrics--that clarifies interpretation context for each metric class. Third, it provides a diagnostic analysis of CPI vocabulary mismatch, quantifying how training-to-evaluation annotation overlap inflates enrichment metrics and contributes to the observed mechanism-clustering gap.

Held-out perturbation metrics (genetic perturbation PCC=0.442; novel drug PCC=0.3047; directional accuracy=73.8% on top-5% effect-size genes) characterize platform prediction capability. CPI enrichment was computed on the Touchstone CPI evaluation set. As shown in Figure 2, the initial EF of 139 at top 0.5% is inflated by a 54.9% overlap with ChEMBL-derived training CPI pairs, dropping to 109 when this overlap is removed (the LINCS mechanism-annotated compound-target validation set; 797 compounds × 225 targets; an approximation, not a strict hold-out); because 54.9% of its positive pairs overlap with our ChEMBL-derived training CPI pairs (the EF drops to 109 after removing overlapping pairs), this metric primarily reflects ranking consistency within the training-derived annotation space rather than out-of-distribution generalization; we report it here for reference only. The self-consistency Mantel rho (0.852) was computed on Subramanian PCL [8] mechanism annotations (filtered to >= 3 drugs per class: 48 classes, 366 compounds), confirming the model retains compound mechanism structure internally. Values obtained on different PCL sets should not be directly compared. All metrics should be interpreted within their respective evaluation contexts.

**Figure 2.**
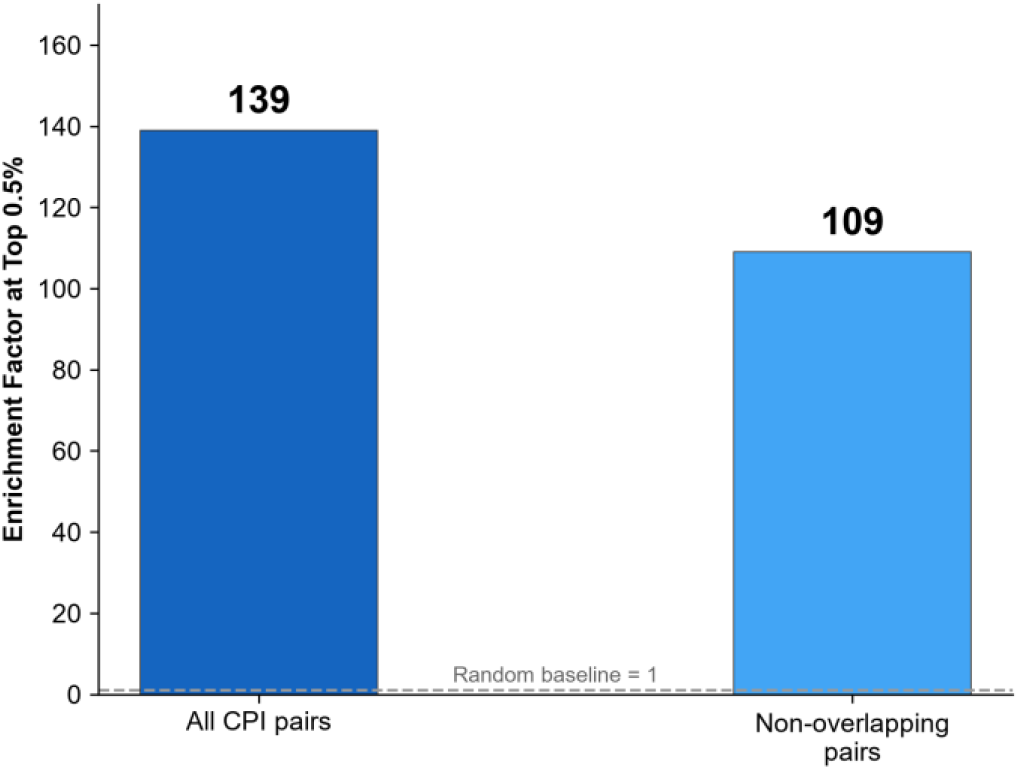
CPI enrichment factor is inflated by training set overlap. EF at top 0.5% for all CPI pairs (139) vs non-overlapping pairs after removing 54.9% overlap with training data (109). Dashed line: random baseline (EF=1).

### 4.2 Rationale for Static Protein Embedding Anchoring

For each of the 4,994 perturbable genes, the protein sequence was taken from the human reviewed (Swiss-Prot) UniProt proteome (reviewed:true, organism: 9606), matched by gene symbol from the FASTA header (GN= field); 12 genes without a matched protein sequence were assigned zero vectors. ESM2-8M [13] encoding was performed once during preprocessing on an NVIDIA RTX 3050 Ti GPU (4 GB VRAM): per-residue representations from the top layer were mean-pooled to 320-dim vectors, which anchor the first 320 dimensions of the 512-dim gene embedding table. The gene embedding table is frozen during finetuning and subsequent training. Our simplified design deliberately isolates the effect of CPI alignment on the compound embedding space through a static protein prior, but as a consequence protein structural information is not integrated into the prediction pathway. This constitutes a key simplification relative to the original trainable protein GNN, and may contribute to the observed SMD gap. Incorporating MSA and trainable GNN encoding is a direction for future work.

### 4.3 SMD Gap Analysis

Because SMD was computed on the same MoA evaluation table with the same per-class filtering criterion (>25 drugs per MoA class) as the original study (Table 3 notes), we present the comparison as a reference; differences in training configuration (CPI pairs, protein encoder) preclude a strict reproduction comparison: the mechanism-clustering SMD of the learned embedding is 1.636, exceeding the ECFP4 Tanimoto fingerprint baseline (1.613) but remaining below the original study’s 1.85 (88.5% attainment). We hypothesize two factors for the remaining gap: (1) CPI vocabulary mismatch: the CPI training pairs used here are based on ChEMBL annotations for the LINCS BRD compound vocabulary (19,161 unique pairs); these differ substantially from the compound–target pairs used in the original study, and may map some structurally dissimilar compounds to similar vectors based on noisy target annotations. (2) Objective conflict: minimizing transcriptional MSE may separate compounds with different effects even if they share a mechanism. Alternative explanations (the simplified protein encoder, frozen gene encoder limiting contrastive learning, hyperparameter sensitivity, data-distribution differences) remain plausible and should be systematically tested in future ablation studies.

### 4.4 Limitations

(1) Single-gene direction prediction remains noisy (60.8% genome-wide); the platform is a candidate-screening and hypothesis-generation tool, not a substitute for experimental validation. (2) The protein encoder is simplified relative to the original. (3) CPI training data has reached its physical ceiling under the current compound vocabulary. (4) The SMD evaluation dataset reflects properties of well-characterized therapeutics and is not representative of screening-library compounds. (5) The original study released code and pretrained models [6,7]; our platform was built from the paper description and did not directly use their released training pipeline, so implementation differences may remain. (6) The increased cell-line coverage introduces greater biological heterogeneity, which itself can lower aggregate performance metrics; genome-wide output (12,328 genes) is more challenging than the original landmark-gene prediction. The 192 randomly initialized and frozen dimensions in the gene embedding table may contribute to suboptimal gene representation; these dimensions were inherited from an earlier 512-dim architecture and retained for checkpoint compatibility, and we did not ablate this design choice. Future work could explore full-dimensional ESM2 initialization or trainable gene embeddings. Future work includes: ablation studies to isolate the contribution of each architectural component; incorporating the original MSA and protein-similarity GNN pipeline; extending to single-cell resolution; and building domain-specific CPI annotation datasets that match the compound vocabulary.

## 5. Conclusion

This study characterizes the scalability of perturbation-to-phenotype prediction at an unprecedented scale of 162 cell lines, 32,039 compounds, and genome-wide output. Three key findings emerge: (1) per-cell-line prediction fidelity is strongly cell-type-dependent, with normal/primary lines achieving mean PCC=0.480 vs. 0.296 for cancer lines, identifying cancer heterogeneity as a primary scalability bottleneck; (2) mechanism-clustering SMD remains moderately stable during scaling (1.636 vs. original 1.85; 88.5% retention at 32x coverage), but CPI vocabulary mismatch between ChEMBL-derived training pairs and the evaluation compound set limits further improvement; and (3) genome-wide directional accuracy on high-effect genes reaches 73.8%, but per-gene recovery is pathway-dependent, as illustrated by the mixed success of case queries across glucocorticoid, proteotoxic stress, adipogenic, and metabolic pathways. These findings generate testable hypotheses about the sources of model failure, and provide a diagnostic framework for identifying where and why virtual cell models degrade--informing systematic ablation studies targeting the CPI vocabulary gap, protein encoder design, and cell-type-aware training strategies.

## Data and Code Availability

All training data are from public repositories: LINCS L1000 (GSE92742, level 5 moderated Z-scores), DepMap CRISPR (23Q2, CERES scores), ChEMBL (release 37), and GEO perturbation datasets (GSE61302, GSE11917, GSE17579, GSE20986, GSE21413, GSE25941, GSE33622, GSE50378, GSE50397, GSE53751, GSE62914, GSE22886, GSE60235). Preprocessed data matrices and trained model weights (g2cp_full_cpi_v7.pt) are available upon reasonable request. Source code is publicly available on GitHub: https://github.com/wkr112344/G2CP-virtual-cell. An interactive API and Web frontend are included in the repository.

## Supplementary Materials

Supplementary Table S1: Per-cell-line held-out drug perturbation PCC values for all 120 cell lines (CSV format). Supplementary File S2: Model checkpoint (g2cp_full_cpi_v7.pt) and evaluation scripts for held-out prediction, directional accuracy, mechanism-clustering SMD, Mantel rho, and CPI enrichment (Python, PyTorch). Supplementary File S3: Interactive query API (Flask, port 8766) for on-demand perturbation-to-phenotype predictions. All supplementary files are available in the same public repository as the code and training data.

## Declarations

## Ethics statement

This study used exclusively publicly available, anonymized datasets and involved no human or animal subjects; therefore, no ethical approval was required.

## Author contributions

K.W. conceived the study, implemented the core architecture and training pipeline, conducted all computational experiments, and wrote the manuscript. L.Z. developed the interactive web interface and visualization components. C.Q. contributed to evaluation benchmarking and biological validation data curation. All authors reviewed and approved the manuscript.

